# Label-free screening of brain tissue myelin content using phase imaging with computational specificity (PICS)

**DOI:** 10.1101/2021.03.22.436473

**Authors:** Michael Fanous, Chuqiao Shi, Megan P. Caputo, Laurie A. Rund, Rodney W. Johnson, Tapas Das, Matthew J. Kuchan, Nahil Sobh, Gabriel Popescu

## Abstract

Inadequate myelination in the central nervous system is associated with neurodevelopmental complications. Thus, quantitative, high spatial resolution measurements of myelin levels are highly desirable. We used spatial light interference microcopy (SLIM), a highly sensitive quantitative phase imaging (QPI) technique, to correlate the dry mass content of myelin in piglet brain tissue with dietary changes and gestational size. We combined SLIM micrographs with an AI classifying model that allows us to discern subtle disparities in myelin distributions with high accuracy. This concept of combining QPI label-free data with AI for the purpose of extracting molecular specificity has recently been introduced by our laboratory as phase imaging with computational specificity (PICS). Training on nine thousand SLIM images of piglet brain tissue with the 71-layer transfer learning model Xception, we created a two-parameter classification to differentiate gestational size and diet type with an accuracy of 82% and 80%, respectively. To our knowledge, this type of evaluation is impossible to perform by an expert pathologist or other techniques.

## I. INTRODUCTION

Myelin is a proteolipid-rich membrane that covers axons and provides the necessary insulation to effectively transmit electrical neural signals throughout various brain regions ^1^. Myelination of fiber bundles is one of the longest brain maturation processes in humans ^2^. Proper myelin development during the perinatal period is crucial for network integration and higher brain functioning ^3,4^ and remains vital in adulthood ^5^. The rapid growth interval during the perinatal period is a decisive time for neural development and is an especially important stage for infants of small gestational age (SGA). Intrauterine growth-restricted (IUGR) and low birth weight (LBW) infants are particularly affected by insufficient myelination. Newborns with such deficiencies are at greater risk of morbidity and mortality ^6^ and display problematic neurological effects that include learning impairments, behavioral problems, neuropsychiatric irregularities, and seizure disorders ^7,8^. The development and analysis of dietary treatments designed to minimize the cognitive issues correlated with IUGR and LB are therefore of great importance.

Different techniques have been used previously for assessing myelin density in biological samples. Luxol Fast Blue (LFB) is a dye that stains myelin blue in tissue fixed with formalin ^9^. In terms of its spatial distribution, LFB provides information on the presence of myelin, but does not allow for its direct quantification. Magnetic resonance imaging (MRI) enables *in vivo* visualization of human brain structures ^10^ and provides a description of myelin concentrations ^11^. However, detection of myelin with MRI is implicit, relying on water proton spins. Although there has been evidence of a decent correspondence between MRI and LFB staining, MRI remains a low sensitivity method ^12^. Proton induced X-ray emission (PIXE) provides a semi-quantitative determination of myelin components within a sample, through phosphorous concentrations, but has low resolution and requires complex and expensive equipment ^10^.

Quantitative phase imaging (QPI) ^13–32^ is a label-free imaging approach that can evaluate pathlength changes in biological samples at the nanometer scale. QPI has numerous medical diagnostic applications ^33^. Di Caprio et al have applied QPI to research sperm morphology ^34^, and Marquet et al. have used QPI to study living neurons ^35^, Lee et al. to study cell pathophysiology ^36^ and Din et al. to examine macrophages and hepatocytes^37^. Conventional quantitative phase methods, however, use coherent light sources that tarnish image contrast with speckles. With the use of a broadband field, phase map. We showed that appropriate for gestational age (AGA) piglets have increased internal capsule myelination (ICM) compared to small for gestational age (SGA) piglets, and that a hydrolyzed fat diet improves ICM in both^41^. However, this analysis was largely manual.

Recently, there has been growing interest in applying the capacity of AI to investigate specific datasets in medical fields ^42–49^. AI has special image processing capabilities to discern multi-faceted features that would otherwise elude trained pathologists. Deep convolution networks provide the opportunity to test thousands of image related feature sets to recognize specific tissue configurations ^50,51^.

Here, we apply phase imaging with computational specificity (PICS) ^52–54^ a new microscopy technique that combines AI computation with quantitative data to extract precise molecular information. Specifically, we combine deep learning networks with SLIM data to define subtle myelin SLIM overcomes this disadvantage, and measures nanoscale information and dynamics in live cells by interferometry ^38^.

We have previously analyzed piglet brain tissue using color spatial light interference microscopy (cSLIM) ^39,40^, which uses a brightfield objective and an RGB camera, and generates 4 intensity images, one of which is a standard LFB color image. Thus, cSLIM simultaneously yields both a brightfield image and a variations in brain tissue, a strategy undertaken for the first time to our knowledge. We used a SLIM-based tissue scanner in conjunction with deep learning methods to classify the associated gestational size and diet of the tissue, which is inherently linked to myelin distribution and mass density. Such a system does not require staining of tissue. However, we performed our measurements on LFB stained samples and computationally normalized the phase maps to account for the effects of the stain ^39^.

## II. METHODS

### A. Brain tissue samples

Tissues were derived from piglets as described in [16]. Associated diets and gestational sizes, as well as tissue slide preparations, are fully described in ^40^. Briefly, piglets were acquired at 2 days of age from the University of Illinois Swine Farm and underwent limited farm processing. SGA was defined as piglets weighing 0.5-0.9 kg at birth, and piglets weighing 1.2-1.8 kg at birth were classified as AGA. Under standard conditions, as defined in a previous publication ^55^, piglets were individually placed in a caging system and randomly assigned to hydrolyzed fats (HF) or control (CON) diet treatment groups in an arrangement of size (AGA or SGA) and diet (CON or HF). The final extracted brain tissues were cut into 4 μm thick sections, mounted on glass slices and subsequently stained with LFB. An example of a coronal section is shown in Figure 1A. All animal care and experimental procedures were approved by the University of Illinois at Urbana-Champaign Institutional Animal Care and Use Committee, in accordance with the National Research Council Guide for the Care and Use of Laboratory Animals.

**FIG. 1.**
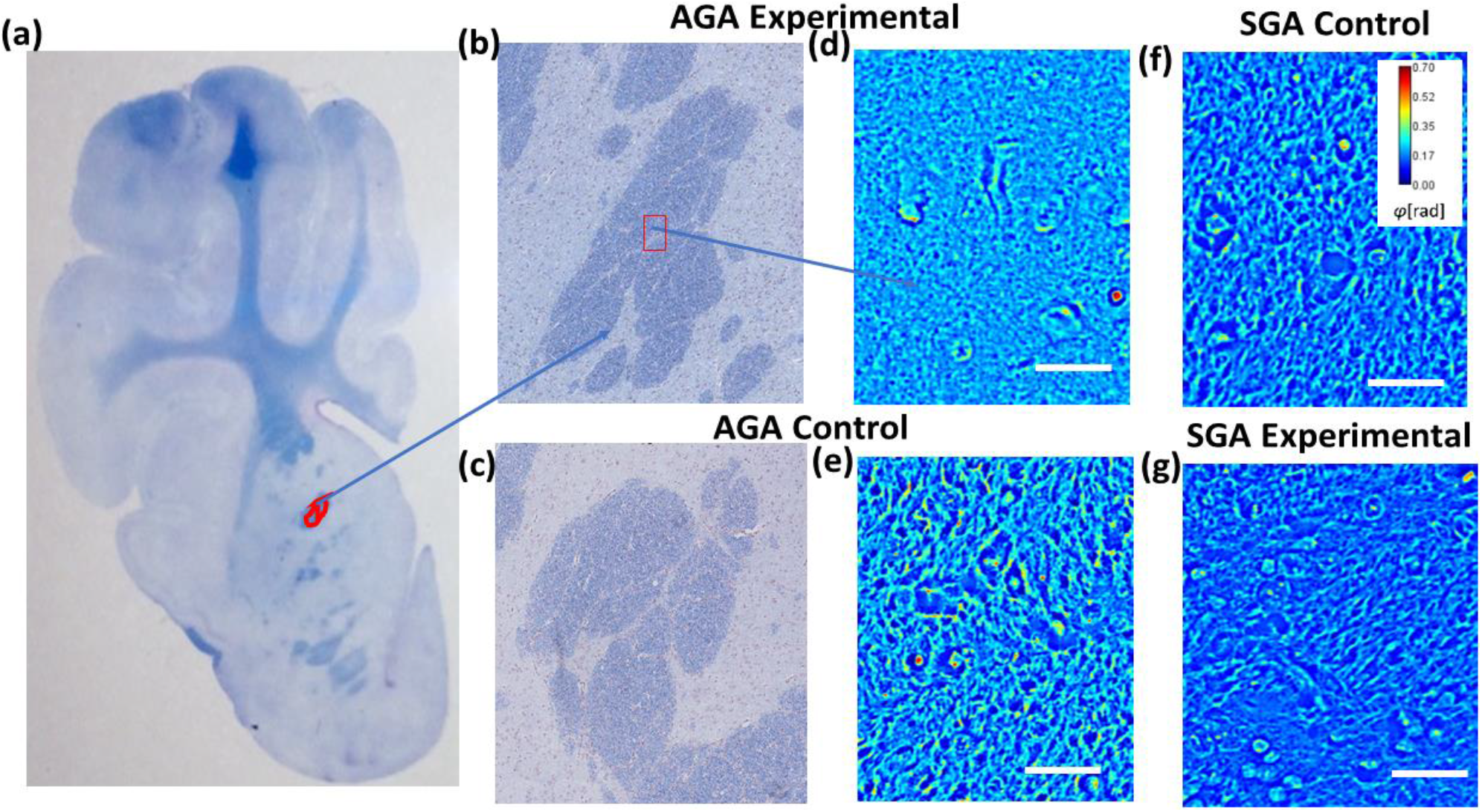
(a) Image of one of the 16 brain sections with the internal capsule demarcated in red. (b) Stitched mosaic of an internal capsule of an AGA piglet with experimental diet obtained using 625 cSLIM images. (c) Stitched mosaic of an internal capsule of an AGA piglet with control diet obtained using 625 cSLIM images. (d) Sample AGA experimental-diet frame, (e) sample AGA control-diet frame, (f) sample SGA control-diet frame, (g) sample SGA experimental-diet frame. Scale bar 50 μm.

### B. Phase Imaging with Computational Specificity (PICS)

We have combined deep learning with SLIM data to predict gestational size and diet regimen from single images. Our label-free SLIM scanner comprises custom hardware and in-house developed software. The SLIM principle of operation relies on phase shifting interferometry applied to a phase contrast setup (see Ref. 30 for details). We shift the phase delay between the incident and scattered field in increments of π/2 and acquire 4 respective intensity images, which suffices to extract the phase image unambiguously. Figures 1D-G show examples of SLIM images of piglet brain tissue corresponding to the area of the internal capsule (IC).

### C. Deep learning model

We employed a transfer learning approach in our deep learning framework to construct our machine learning classifier. This technique is recommended for training a model when there is a relatively small number of image instances.

As outlined in Figure 2A, we selected the Xception model, which comprises 71 layers and has been pretrained on a large dataset of over 1.6 million images of different sizes and groups. Due to its robust feature extraction capacity and superior performance with our data instances, this model was selected over alternatives such as ResNet ^56^ and MofileNet ^57^. We finetuned the base model to include two dropout layers of 0.75 (Fig. 2. B).

**FIG. 2.**
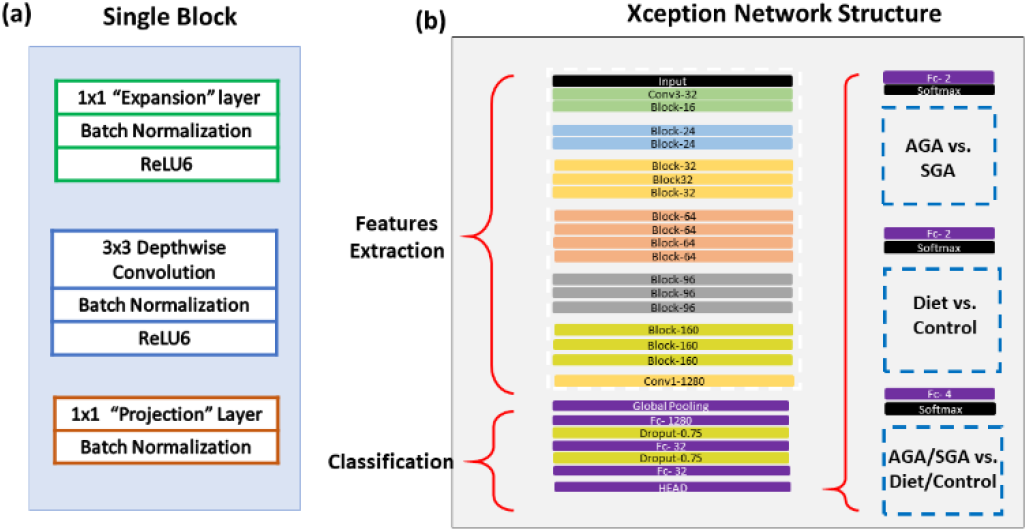
(a) Machine learning Xception network components of a single block. (b) Entire network structure with a finetuned classification segment and three different output models.

## III. RESULTS

### A. Data

Our training data included 9008 SLIM images of piglet brain samples, obtained from 16 sections of different piglets divided equally between four categories: AGA-diet, SGA-diet, AGA-control, and SGA-control. Full IC reconstructions and sample frames are illustrated in Figures 1B-C. The phase distributions of the various groups are displayed in Figures 3A-F. Each graph displays one of the six possible pairings of the four categories. Figures 3A-B show the different phase distributions caused by diets in the same gestational classes, while figures 3C-D show such differences caused by gestational age in the same diet categories. Figures 3E-F contrast mixed diet-age distributions. The closeness and extent of overlap for each combination illustrates the minute numerical differences in the images themselves, suggesting that diagnostic capabilities are largely due to the differences in the spatial distributions. This is further substantiated in the statistical differences in dry mass measurements of the same samples found in our previous study ^40^ only after applying binary myelin masks to the quantitative data.

**FIG. 3.**
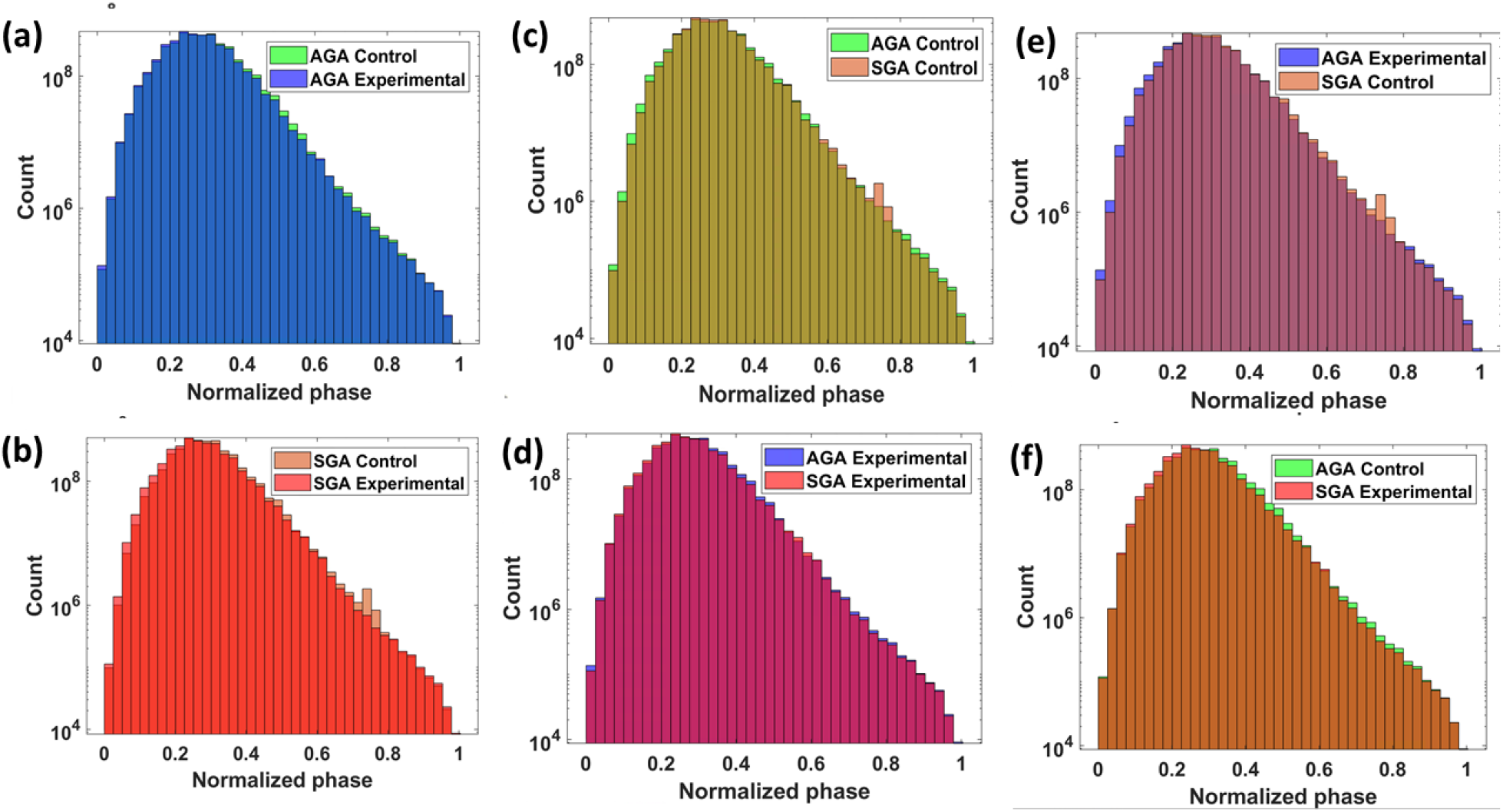
Histograms of the count of pixel data in SLIM images for (a) AGA groups, (b) SGA groups, (c) Control diet groups, (d) Experimental diet groups, and (e-f) mixed gestational age and diet groups.

### B. Model accuracy and loss

Model accuracy and losses for three types of classifications are shown in Figure 4. In the case of distinguishing brain tissue based on diet regimens, we obtained an accuracy of 80% (Fig. 4 A.) and a loss of 0.35 (Fig. 4. B). These results are significant considering that the subtlety of these difference would otherwise be undiscernible to a trained histopathologist. There is negligible underfitting or overfitting in these models and they could be characterized as having appropriate and balanced fitting. In the case of classifying phase maps based on gestational size, the results were slightly stronger with an accuracy of 82% (Fig. 4. C.) and a loss under 0.3 (Fig. 4. D). There is minimal underfitting in this model, however the training loss is noisy near the final epochs, likely due to the large number of parameters being evaluated with all the weights set to true. Lastly, the results for the classification of both diet and gestational age categories are, as is expected, considerably lower with an accuracy of 63% and a slight degree of underfitting (Fig. 4. E). The loss values are also higher than individual comparisons, tapering off around 0.8 (Fig. 4. F).

**FIG. 4.**
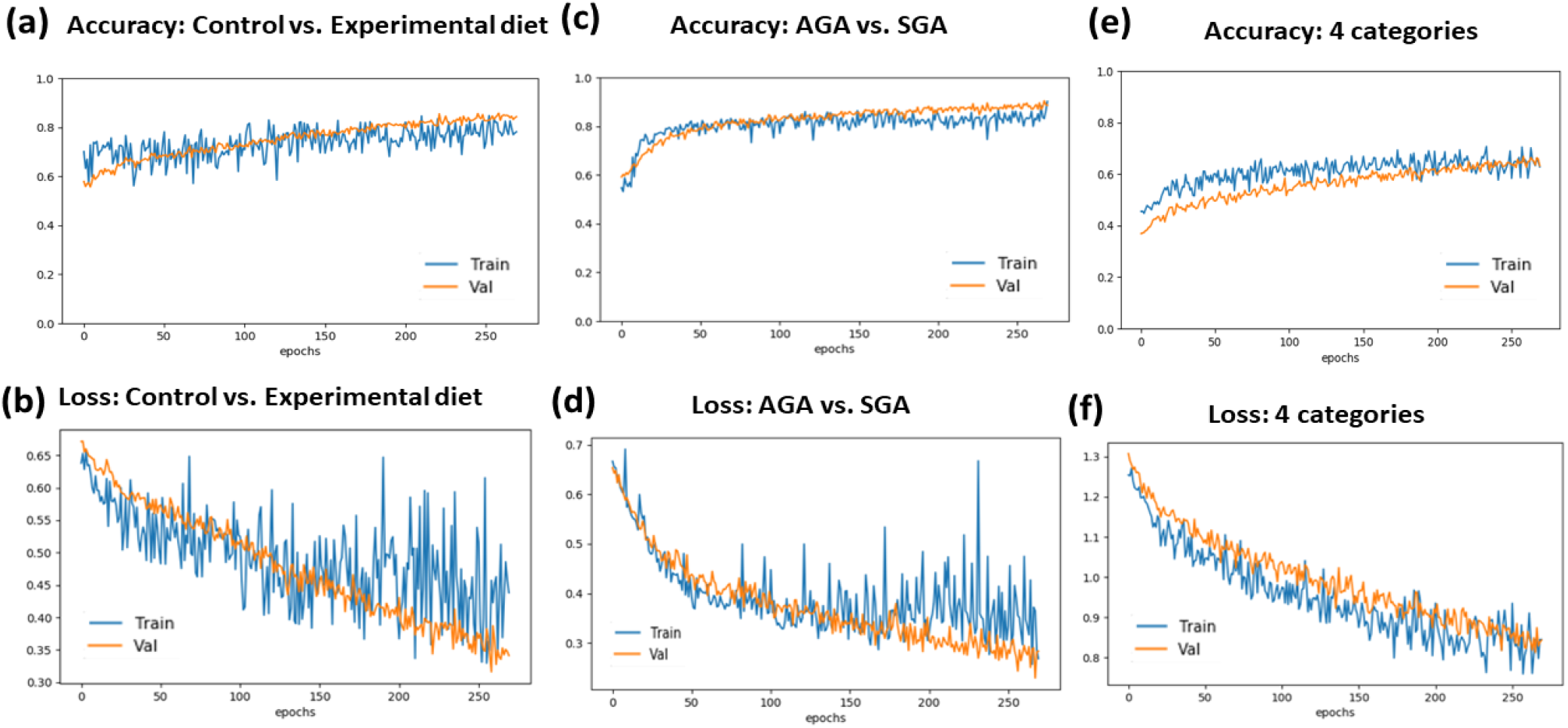
Plots for diet classification (a) accuracy and (b) loss, (c) gestational size accuracy and (d) loss, and (e) all categories accuracy and (f) loss.

### C. Confusion matrices for validation and loss

The confusion matrix offers a quantitative indication of the performance of a classifier. There are four classes in our confusion matrix: “Diet - AGA”, “Control - AGA”, “Diet - SGA”, and “Control - SGA”. This confusion matrix can have three kinds of errors: the sample can be labelled incorrectly in terms of diet, gestational size, or both. In the case of a perfect classification model, the confusion matrix is diagonal with only true negatives or true positives. Figure 5A shows a 4×4 confusion matrix for the classification of each category on all test images from 16 slides.

**FIG. 5.**
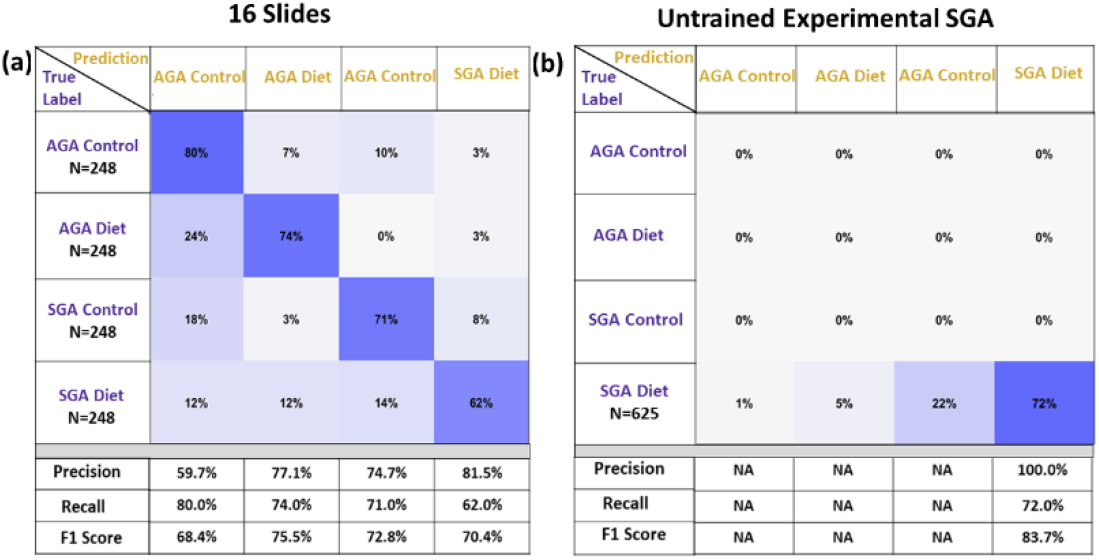
(a) Confusion matrix for classification of all four categories on the 16 slides that were used for training, and (b) confusion matrix of an untrained Experimental diet-SGA slide.

The first row, corresponding to the Control-AGA category, indicates 80% of samples labelled correctly with most errors attributed to designating the samples with the same diet but with a *small* gestational size. The second row, corresponding to Diet-AGA, has 74% correct labelling, with most errors due to a mismatch of the diet type. The third row, for Control-SGA, has 71% correct labeling and the last row, for Diet-SGA, has only 62% correct labelling, with 14% mislabeled as Control-SGA. The AGA categories outperform those of SGA, presumably due to a lower myelin abundance inherent in the smaller gestational size, which may have been counterbalanced by the experimental diet, thereby confusing the Diet-SGA category with either AGA classes.

In order to validate our model on slides that were not used for training, we evaluated images from an untrained slide that is associated with an experimental diet and SGA. The results, shown in a confusion matrix (Fig. 5.B), were better than anticipated, exceeding the performance of test images in this category, with a true positive rate of 72 % for both size and diet, with 75% for just diet, and 95% for just size.

## IV. CONCLUSIONS

Current histopathological findings depend on manual investigations of stained tissue slices under a microscope by a trained pathologist. The alternative methods of assessing myelin density, such as MRI and PIXE, are indirect, cumbersome, and costly. Here, we present evidence that our method of combining AI with spatial light interference microscopy (SLIM) can quickly determine differences in myelin content without the use of molecular stains or manual analysis. This is an important contribution to neuroscience, especially given the significance of myelination in brain development and the current challenges of measuring myelin quantitatively.

We demonstrated that applying AI to SLIM images delivers excellent performance in classifying single phase maps of brain tissue to detect the level of myelin adequacy. The approximately 80 percent accuracy outcomes for both binary distinctions, and 62 percent for all four categories, indicate that the proposed method may be useful in quick screenings for cases of suspected myelin disorders. These results are significant as it would otherwise be impossible for a trained histopathologist to distinguish such myelin discrepancies. Not only does this technique offer automatic screening, but multiple tissue samples can be analyzed rapidly as the overall throughput of the SLIM tissue scanner is comparable with that of commercial whole slide scanners.

Future scope includes evaluating myelin content at the single axon level by creating specificity masks using fluorescent tags for constituent proteins, such as the proteolipid protein. Employing PICS to mimic such tags would facilitate investigations into the dynamic generation of myelin around axons in real-time.

## ACKNOWLEDGEMENTS

This work was supported by the National Institute of General Medical Sciences (NIGMS - https://www.nigms.nih.gov/) grant GM129709, the National Science Foundation (NSFhttps://www.nsf.gov/) grant CBET‐ 0939511 STC, the National Institutes of Health https://www.nih.gov/grant CA238191, as well as Abbott Nutrition C3123.

## DATA AVAILABILITY

The data that support the findings of this study are available from the corresponding author upon reasonable request.

